# Sustained dynamics and modality-specific network reconfigurations define crossmodal prediction error

**DOI:** 10.1101/2025.10.29.685284

**Authors:** Maria Niedernhuber, Federico Raimondo, Jacobo Sitt, Tristan Bekinschtein

## Abstract

Predictive processing theories posit that the brain continuously generates expectations about incoming sensory inputs, updating them through prediction errors. While extensively studied within single modalities, it remains unclear how prediction errors unfold when expectations and violations occur across different senses. Using a local-global hierarchical oddball paradigm combined with high-density EEG in 47 participants, we contrasted unimodal and crossmodal prediction errors across auditory, visual, and somatosensory domains. We found that crossmodal violations elicit temporally sustained cortical responses which diverge from the transient, localised dynamics observed for unimodal prediction errors. Temporal decoding revealed that crossmodal effects maintain shared prolonged neural representations, suggestive of supramodal integration across all levels of cortical processing. Computational modelling further demonstrated that crossmodal prediction errors reorganize effective connectivity within and between sensory hierarchies, engaging distinct early cortical pathways depending on the sensory combination. Our findings refine hierarchical predictive coding for crossmodal transitions by demonstrating that, unlike unimodal prediction errors, crossmodal prediction errors recruit dedicated prolonged supramodal representations and flexibly adapted modality-specific networks.

## 1 Introduction

A key question in neuroscience is how the brain predicts novel events in an uncertain multisensory environment (Clark 2013; Friston 2005; Kok and de Lange 2015). According to predictive coding theories of brain function, neural signals encode a prediction error which arises from a comparison of predicted and actual sensory information. Prediction errors are transmitted along a hierarchy of cortices ordered from predominantly modality-specific early to supramodal late cortices (Niedernhuber et al. 2022). At every level of the cortical hierarchy, the prediction error is forwarded to the subsequent level and updated with top-down predictions until the prediction error is minimal (Friston 2008; Rao and Ballard 1999; Lee and Mumford 2003). To enable multisensory predictions, the cortex needs to integrate information from different senses presented in the same spatiotemporal context (Talsma 2015; Cao et al. 2019). Crossmodal prediction errors can be detected when sensory expectations are generated and violated in different sensory modalities (Weilnhammer et al. 2018; Huang et al. 2023; Keil and Senkowski 2018; Roa Romero et al. 2016). There is a current of experiments refuting classical theories according to which early sensory cortices are strictly modality-specific (Rao and Ballard 1999). These experiments demonstrate that multisensory integration is pervasive across the cortical hierarchy, albeit not to the same extent (Driver and Noesselt 2008; Macaluso and Driver 2005; Ghazanfar and Schroeder 2006; Foxe and Schroeder 2005, 2005). Accordingly, neural signatures of crossmodal prediction errors are present in primary (Matusz, Retsa, and Murray 2016) as well as higher-order cortices (Sabio-Albert, Fuentemilla, and Pérez-Bellido 2025).

Neural responses to sensory irregularities such as the Mismatch Negativity (MMN) can be used to understand how prediction errors are encoded in the brain. The MMN is a negative response elicited 100-200 ms after a sensory irregularity is introduced in a spatiotemporally confined sensory context (Risto Näätänen 1990; R. Näätänen et al. 2001). Originally discovered in the auditory modality (Risto Näätänen, Gaillard, and Mäntysalo 1978), the MMN can also be elicited using tactile and visual paradigms Cammann (1990). Numerous studies located the neural generators underlying the MMN in early modality-specific cortices, such as the primary visual, somatosensory and auditory cortex, as well as the inferior frontal gyri (Grundei et al. 2023; Chennu et al. 2016; Garrido et al. 2009). Aligning with the idea of a cortical hierarchy with increasingly supramodal levels, our group and others demonstrated a supramodal contribution to the MMN likely originating in the prefrontal cortex (Grundei et al. 2023; Niedernhuber et al. 2022). Although the MMN is typically studied within a sensory modality (Risto Näätänen 1990; R. Näätänen et al. 2001; Maekawa et al. 2009; Cammann 1990; Fardo et al. 2017; Gijsen et al. 2021; Naeije et al. 2016; Chennu et al. 2016; Garrido et al. 2009), there is evidence that crossmodal sensory expectation violations elicit a MMN-like crossmodal prediction error signal (Huang et al. 2024; Ghosh, Talwar, and Banerjee 2024). Complementarily, Grundei et al. (2023) used a trimodal roving oddball design in humans to show that crossmodal deviance engages a shared supramodal network including temporoparietal and inferior frontal regions (Grundei et al. 2023).

In this study, we used unimodal and crossmodal version of the local-global hierarchical oddball paradigm to contrast unimodal and crossmodal prediction errors (Bekinschtein et al. 2009; Huang et al. 2024). The local-global oddball paradigm introduces hierarchically nested sensory expectation violations in the visual, auditory and somatosensory modality. Sensory irregularities can occur at a local level (deviant stimulus at the end of a trial) or a global level (deviant trials within a block). In our version of the paradigm, predictions of local sensory violations were either generated and violated in different sensory modalities, or the same sensory modality. We hypothesised that local sensory expectations generated and violated in different sensory modalities produce a crossmodal local effect. We also examined whether crossmodal local effects might be supported by a common sustained response at a higher level of the cortical hierarchy. Finally, we hypothesised that inferior frontal cortical hubs might consistently be involved in crossmodal prediction error routing, in addition to other regions unique to each crossmodal contrast (Ghosh, Talwar, and Banerjee 2024). To test this hypothesis, we compared effective connectivity architectures between unimodal and crossmodal local effects to test this hypothesis using Dynamic Causal Modelling (DCM) and Parametric Empirical Bayes (PEB).

## 2 Methods

### 2.1 Participants

54 individuals (35 female, 19 male, mean(±STD) age: 25.20(±4.1) years) participated in the experiment. We only asked people aged 18-80 without neurological and psychiatric conditions or auditory, visual or tactile impairments. We excluded 7 participants due to a recording error. Recruitment was performed using the RISC system at the Centre National de la Recherche Scientifique (CNRS) in France. Participants gave their informed consent, and received €40 in compensation.

### 2.2 Paradigm

We designed crossmodal and unimodal version of the local-global oddball paradigm (Bekinschtein et al. 2009), in which sensory irregularities are presented locally (deviant stimulus within a trial) or globally (deviant trials within a block). Details are shown in Figure 1). Trials either consisted of five identical stimuli (locally standard trials), or four identical stimuli followed by a deviant (locally deviant trials). In a block, repetitions of a trial type (globally standard trials, 80% of trials in a block) are occasionally interrupted by the other trial type (globally deviant trials, 20% of trials in a block). In block type XX, locally standard trials were interrupted by locally deviant trials, and in block type XY vice versa. Standard stimuli were presented ipsilaterally (ear, visual hemifield, or wrist), followed by a contralateral deviant stimulus. 50% of trials in a block started with left-sided stimulation (and vice versa). 31 trials in each block were globally deviant. Globally deviant trials were preceded by 3, 4, or 5 globally standard trials. 24 globally standard trials were presented at the start of each block to induce an expectation of a global regularity.

**Figure 1:**
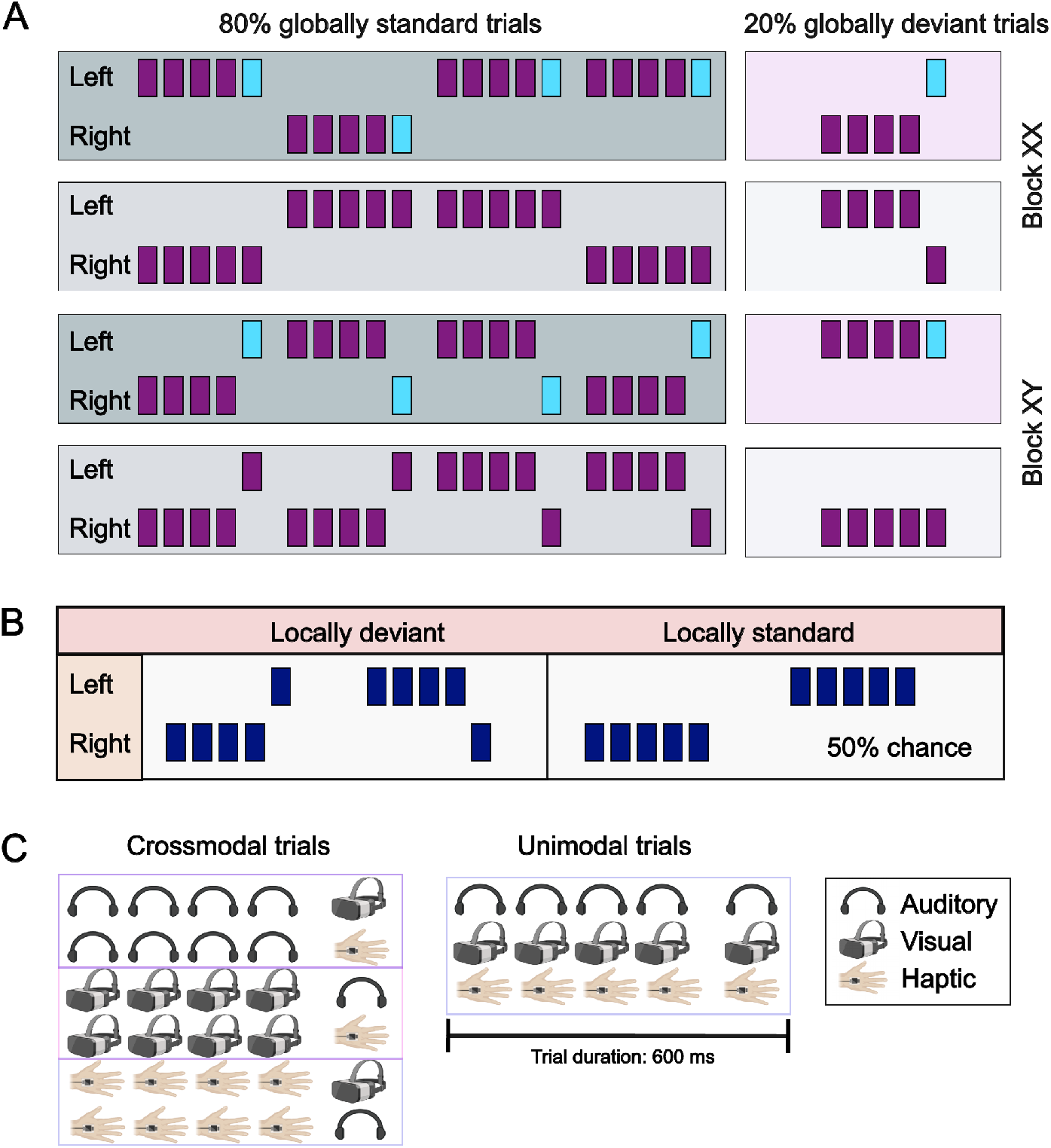
Paradigm. Block design with 80% globally deviant and 20% globally standard trials in each block. In block XX, locally standard trials are also globally standard, whereas in block XY they are globally deviant (and vice versa for locally deviant trials). Colors (purple/blue) denote the sensory modality in which a stimulus was delivered (A). Locally deviant trials consist of four ipsilateral standard stimuli followed by a contralateral stimulus. Conversely, standard trials consisted of five ipsilateral stimuli. Standard stimuli could randomly start on the left or right side (B). Crossmodal trials consisted of four stimuli in the same sensory modality followed by a stimulus presented in another sensory modality. Unimodal trials consist of five stimuli presented in the same sensory modality (C).

Stimuli in unimodal blocks were sampled from the same sensory modality. Trials in crossmodal blocks were always composed of four ipsilateral unimodal stimuli followed by a contralateral stimulus sampled from a different sensory modality. As a result, there are 9 crossmodal trial types: auditory-somatosensory, somatosensory-auditory, auditory-visual, visual-auditory, visual-somatosensory, and somatosensory-visual. The experiment was performed in three sessions which lasted 4.5 min (20 min total duration including breaks). Each block type was presented twice per session in alternating order (XX-XY-XX-XY). Participants were assigned to one of the following session combinations: 1. auditory-somatosensory, visual-somatosensory, somatosensory (16 participants); 2. visual-auditory, somatosensory-auditory (15 participants), auditory; 3. auditory-visual, somatosensory-visual, visual (16 participants).

Our task was designed to extract a series of cortical markers located at successive hierarchical levels in the cortex: 1. Unimodal local effects manifest as a relative enhancement of amplitude in a time window between 50-250 ms when locally standard trials (i.e. groups of five identical unimodal ipsilateral stimuli) are contrasted with locally deviant trials (i.e. groups of four identical ipsilateral stimuli followed by a contralateral input). 2. Crossmodal local effects can be detected when ERP amplitude time courses of unisensory unilateral groups of five stimuli are compared with groups of four unimodal stimuli followed by an ipsilateral stimulus sampled from a different sense in a time window between 50-350 ms. 3. Global effects were established by contrasting ERP time courses extracted from globally deviant vs standard trials in a time window from 250-600 ms after onset of the last stimulus in a trial in crossmodal and unimodal blocks.

### 2.3 EEG preprocessing

We collected EEG data using a high-density EEG net with 256 channels and a Net Amps 300 amplifier (Electrical Geodesics) at the ICM in Paris. Preprocessing was performed in Matlab 2019b using EEGLAB (based on a pipeline used in (Chennu et al. 2013)). EEG data were resampled from 500 Hz to 250 Hz. 81 electrodes placed around the cheek and neck were categorically removed due to muscle artefacts. We removed 24 globally standard trials which were presented at the start of each block to generate an expectation of the global sensory pattern. We filtered the EEG data between 0.5 and 30Hz with a butterworth filter, and epoched relative to trial onset. Baseline removal was performed in a window of 100 ms before trial onset. We rejected trials with a variance of >350 and channels with a variance of >500 using a semi-automated procedure. Then, we performed independent component analysis and rejected components representing noise using a semi-automated pipeline. Finally, we removed any remaining noise using trial-wise interpolation.

### 2.4 Cluster-based permutation test

For each condition pair, we tested whether deviant trials produce a larger response than standard trials. We compared ERP voltage time series between deviant and standard trials using a cluster-based permutation test implemented in Fieldtrip (Maris and Oostenveld 2007). For each time-channel sample, we ran a two-sided t-test and thresholded t-values at cluster *α* < 0.05. We calculated maximum cluster t by summarising t-values in spatiotemporally adjacent samples and selecting the maximum resulting t-value. Finally, we ran a Monte Carlo algorithm with 3000 random partitions to obtain the Monte Carlo significance probability *α* < 0.05.

### 2.5 Temporal generalisation analysis

To characterize the dynamics of unimodal and crossmodal neural representations, we performed temporal generalization decoding for each participant using MNE-Python (Gramfort et al. 2014). This method assesses whether a neural pattern discovered at a specific moment in time is present at other moments, revealing the stability and dynamics of the underlying representation (King et al. 2014).

For classification, we used a linear logistic regression model with an L2 penalty solved using a Coordinate Descent optimization algorithm in the liblinear library. EEG data were first standardized (z-scored) using a standard scaler at each channel and time point. For each time point in a training set, a classifier was trained to distinguish deviant from standard trials. Due to the difference in trial numbers in both conditions, we adjusted class weights to be inversely proportional to class frequencies. Trained classifiers were tested on their ability to classify trials at every other time point in a separate testing set. We repeated this process for every time point in the training set, resulting in a temporal generalization matrix of classification performance quantified as the Area Under the Receiver Operating Characteristic curve (AUC-ROC). An AUC score of 0.5 indicates chance-level performance. The diagonal of this matrix reflects the instantaneous decoding accuracy, while off-diagonal elements indicate the degree to which a neural pattern is sustained or re-emerges over time.

To characterize neural dynamics within a specific condition (e.g., distinguishing auditory-visual deviant trials from standard trials), we used a 5-fold stratified cross-validation procedure within each participant. We used a hold-out validation approach to test for shared neural representations across different conditions since the training and testing data were drawn from distinct trial sets. A classifier was trained on all trials of one modality (e.g., auditory-visual deviant vs. standard trials) and subsequently tested on all trials from a different modality (e.g., visual deviant vs. standard trials). We performed these procedures for all relevant pairs of modalities within each participant, resulting in a set of within-subject AUC-ROC matrices per condition. To identify decoding performance differences between condition pairs, we performed non-parametric Monte Carlo cluster-based permutation tests with 1024 random partitions (Maris and Oostenveld 2007). We applied one-tailed, one-sample t-tests (*α* <0.05) to assess whether adjacent classification scores were above chance level (AUC = 0.5).

### 2.6 Parametric Empirical Bayes

We used Parametric Empirical Bayes (PEB) to test for differences in effective connectivity between unimodal and crossmodal local effects. Dynamic Causal Modelling (DCM) for ERPs is a Bayesian method to infer effective connectivity between neural sources. DCM are neuronal mass models which describe excitatory and inhibitory dynamics between neuronal subpopulations (Jansen and Rit 1995). Cortical sources are modelled as equivalent current dipoles encompassing excitatory pyramidal cells, spiny stellate cells and inhibitory interneurons. Information flow between sources can be modelled as forward, backward or lateral connections. DCM can also include intrinsic connections which represent the causal influence of a neuronal population over itself. Variational Laplace is used to estimate the strength of the connections between sources in the model. After estimating the probability density of connectivity parameters for each participant, connectivity parameters can be modelled at a group-level using a General Linear Model. PEB refers to the hierarchical linear model constructed over connectivity parameters obtained from subject-wise DCMs. To compare effective connectivity between two conditions at a group level, we can estimate a PEB for each condition and construct a further general linear model over the resulting connectivity parameters (PEB-of-PEBs).

## 3 Results

### 3.1 Crossmodal prediction errors evoke mid-latency neural responses

Using cluster-based permutation testing, we identified crossmodal local effects in a time window between 50-250 ms and crossmodal global effects in a time-window from 300 ms to trial end, regardless of which sensory modalities were converged in the paradigm (Figure 3 and 2). As previously reported (Niedernhuber et al. 2022), the topography of the effects differed between modalities and was characteristic of each stimuli pattern. The auditory local effect was characterized by an initial negative deflection followed by a positivity (t = 160.18, p < 0.001), whereas the somatosensory local effect predominantly exhibited a positive wave (t = 178.76, p < 0.001). In contrast, the visual local effect was marked by a clear negativity (t = 191.91, p < 0.001).

**Figure 2:**
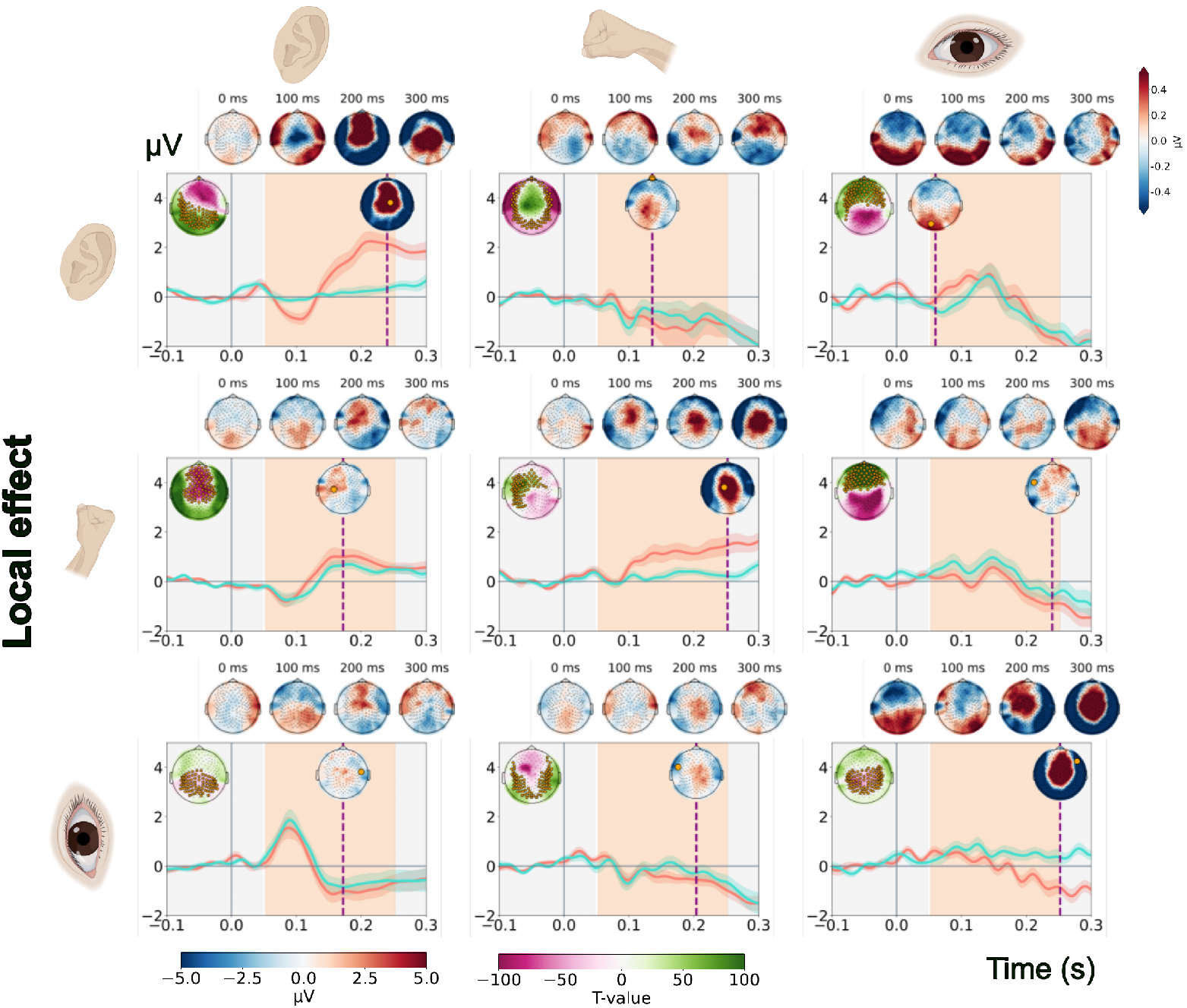
Local effect ERP time courses. ERPs show deviant (salmon) and standard (teal) average time courses with the standard deviation shown shaded obtained at the electrode with the maximum difference between conditions. The point at which both time courses diverge maximally is marked with a purple dotted line and the corresponding topography is shown on top. In the upper left corner, the T-map of the cluster with the largest summarised observed T is shown and electrodes belonging to this cluster are shown in orange. Topographies of the difference between deviants and standards are shown on top of each panel.

**Figure 3:**
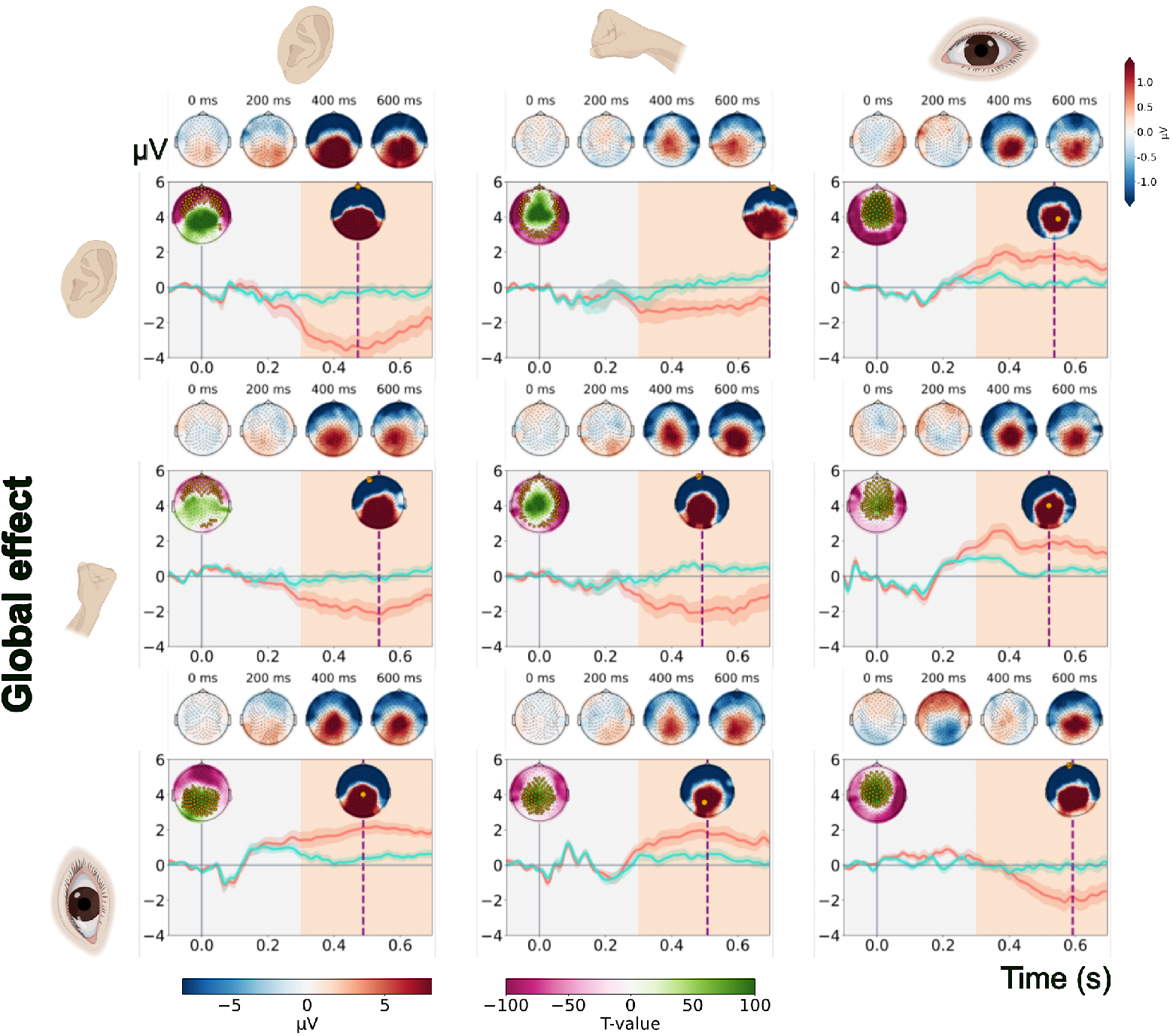
Global effect ERP time courses. ERPs show deviant (salmon) and standard (teal) average time courses with the standard deviation shown shaded obtained at the electrode with the maximum difference between conditions. The point at which both time courses diverge maximally is marked with a purple dotted line and the corresponding topography is shown on top. In the upper left corner, the T-map of the cluster with the largest summarised observed T is shown and electrodes belonging to this cluster are shown in orange. Topographies of the difference between deviants and standards are shown on top of each panel.

Among the crossmodal local interactions, the auditory-visual condition revealed a positive deflection with an early peak occurring before 100 ms (t = 213.85, p < 0.001), while the somatosensory-visual combination demonstrated a later positive peak near 120 ms (t = 198.93, p < 0.001). The visual-somatosensory interaction was associated with a subtle negativity peaking around 200 ms (t = 208.97, p < 0.001), and the auditory-visual pairing showed a positivity followed by a subsequent negativity (t = 193.39, p < 0.001). The somatosensory-auditory condition resulted in a mid-latency positive deflection (t = 149.77, p < 0.001), and the auditory-somatosensory combination similarly elicited a positive wave (t = 196.90, p < 0.001).

Regarding global effects, we observed pronounced negative deflections from approximately 300 ms for the visual (t = 225.55, p < 0.001), somatosensory (t = 181.21, p < 0.001) and auditory (t = 194.86, p < 0.001) conditions. Likewise, large-amplitude negativities were apparent for the somatosensory-auditory (t = 235.59, p < 0.001) and auditory-somatosensory effect (t = 212.57, p < 0.001). In contrast, the visual-somatosensory (t = 168.31, p < 0.001), somatosensory-visual (t = 178.21, p < 0.001), visual-auditory (t = 175.54, p < 0.001) and somatosensory-auditory effects (t = 118.98, p < 0.001) global effects manifested as positive deflections.

### 3.2 Temporal Decoding

Using temporal decoding, we characterised how cortical representations of crossmodal and unimodal prediction errors evolve in time (Figure 4). We also investigated whether, and to which extent, crossmodal and unimodal prediction errors share neural representations.

**Figure 4:**
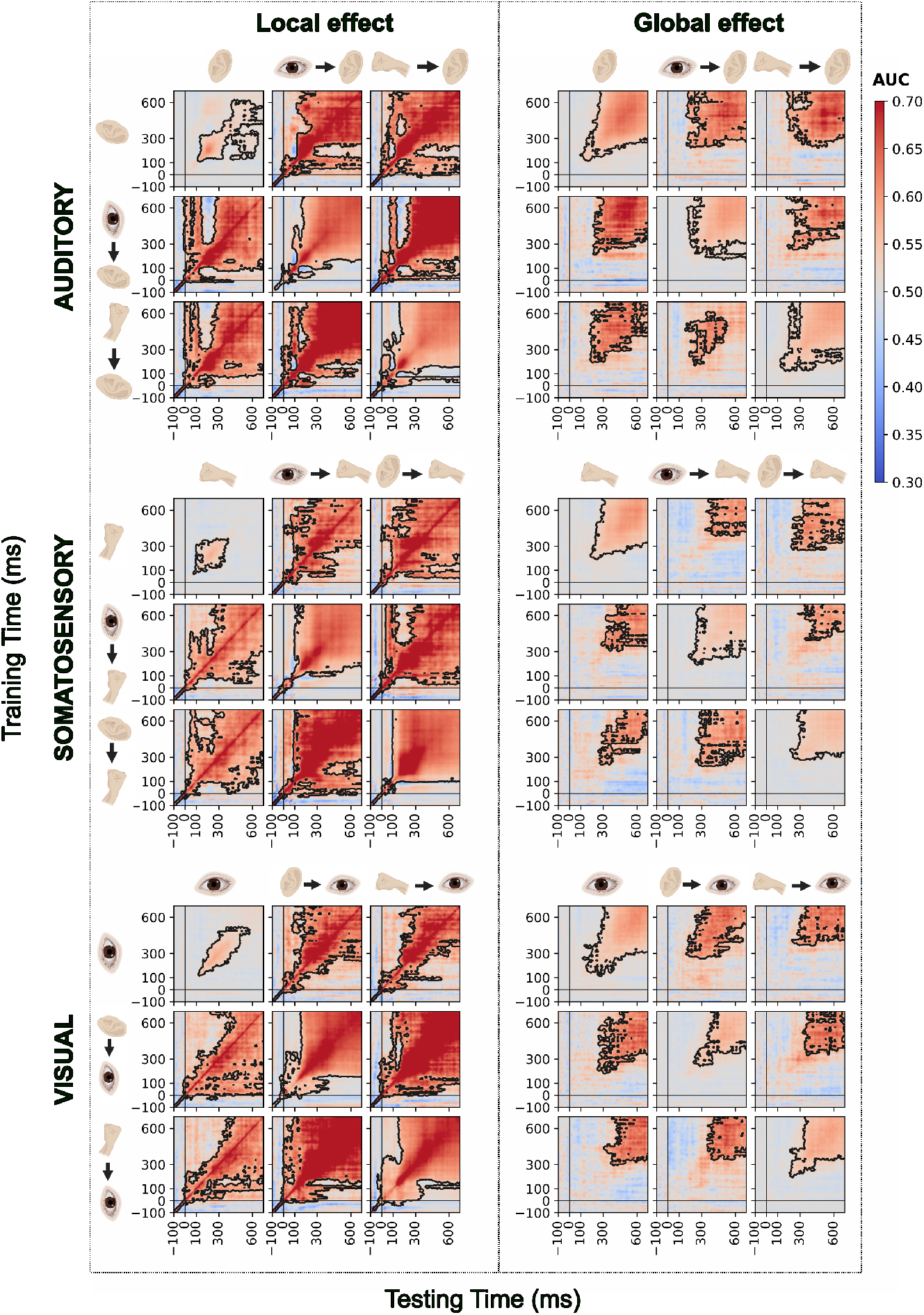
Temporal decoding. Matrices show AUC-ROC scores for distinguishing standard vs. deviant trials for each crossmodal and unimodal local and global contrast. Classifiers were trained at each time point on the y-axis and tested at each time point on the x-axis for local (left) and global (right) effects. In each panel, classification score matrices along the diagonal represent neural dynamics resulting from a comparison of deviant and standard trials within a condition (5-fold cross-validation) whereas the remaining classification score matrices show shared neural dynamics between conditions, resulting from a comparison of deviant vs standard trials (hold-out cross-validation). Rows correspond to the contrast used for training, and columns for testing. Contrasts are indicated with icons (ear: auditory, eye: visual, hand: somatosensory). Single icons represent contrasts within a sensory modality, and icons separated by an arrow contain standard stimuli from the sensory modality on the left followed by a deviant stimulus from the modality on the right. AUC-ROC scores were mapped on a red-to-blue gradient. Black contours mark significant clusters (p < 0.05).

#### Unimodal local effects

Replicating previous findings (King et al. 2014; Niedernhuber et al. 2022), unimodal local effects elicit a mid-latency, short-lived response decodable primarily along the diagonal. For the auditory local effect, temporal decoding revealed short-lived activity between ∼100–300 ms. Likewise, the somatosensory local effect produced a transient activation pattern best decoded from ∼200–350 ms. Finally, we observed briefly maintained activation patterns in a 180–400 ms time window supporting the visual local effect. This suggests that local deviance signals are transient and likely reflect propagation along the cortical hierarchy rather than sustained reverberation within neural circuits.

#### Crossmodal local effects

In stark contrast, crossmodal local effects were maintained across longer temporal windows and demonstrated broader generalisation across time compared to their unimodal counterparts. Local crossmodal violations recruited temporally stable, shared neural representations, though the specific dynamics were dependent on the sensory modality of the deviant stimulus. Auditory crossmodal violations (somatosensory-auditory, visual-auditory) were characterized by a prominent, sustained period of temporal generalization across the entire time window. Auditory crossmodal effects elicited activation along the diagonal which generalised from ∼300 ms until the end of the trial. Similarly, somatosensory local crossmodal effects (auditory-somatosensory, visual-somatosensory) elicited a strong, rectangular pattern of sustained generalization in a mid-latency window, from ∼200 ms until trial end. Finally, visual local crossmodal effects (auditory-visual, somatosensory-visual) initially produced sustained activation for ∼200 ms in an early time window, and then elicited a strong, rectangular pattern of sustained generalization in a mid-latency window, from ∼200 ms until trial end. Taken together, these results show that a multisensory context facilitates sustained, supramodal representations of local prediction errors whose temporal dynamics are shaped by the specific modality being violated. Importantly, we observed temporally extensive shared activation patterns between crossmodal local effects, as well as between crossmodal and unimodal local effects across all conditions. This suggests that crossmodal local effects activate supramodal cortical hubs which sustain a single and stable activation pattern.

#### Unimodal global effects

Temporal decoding of unimodal global effects revealed that they generalise in a late time window with some shifts in onset. As shown in previous work (Niedernhuber et al. 2022), the auditory global effect is maintained from ∼200 ms until trial end, whereas sustained activation started at ∼300 ms for the somatosensory and visual condition.

#### Crossmodal global effects

Auditory (visual-auditory, somatosensory-auditory) and somatosensory (auditory-somatosensory, visual-somatosensory) crossmodal global effects manifested in late generalisation from ∼300 ms. For visual crossmodal effects (auditory-visual, somatosensory visual), generalisation onset was delayed at ∼400 ms. We observed shared supramodal late representations from ∼400 ms until trial end for all crossmodal and unimodal effects. This finding aligns with earlier work from our group suggesting that the global effect is supported by late supramodal activation regardless of sensory modality (Niedernhuber et al. 2022).

### 3.3 Effective connectivity modulations reflect modality-specific adaptation to prediction error context

To characterize the effective connectivity supporting prediction error processing, we first modelled the network architecture for each of the nine local effects, retaining connections with a posterior probability exceeding 0.95 shown in Figure 5. In the three unimodal conditions, a canonical, modality-specific network emerged. This was characterized by excitatory feedforward connections, top-down excitatory input from the IFG to higher sensory areas, and inhibitory feedback from higher to lower sensory cortices. Crucially, in the six crossmodal conditions, this canonical hierarchy of the deviant modality was augmented by significant inter-sensory connections, revealing how the network dynamically integrates the sensory context of the preceding standard stimuli. For instance, when an auditory deviant followed somatosensory standards, the core auditory network was supplemented by an excitatory connection from the right S2 to the right A1 and an inhibitory connection from the right S2 to the right STG. When an auditory deviant followed visual standards, we observed excitatory drive from the right V5 to the right A1. This pattern of cross-hierarchy integration was consistent across all conditions. Visual deviants were influenced by the preceding context via an excitatory connection from the right STG to the right V1 in the auditory-visual condition, and an excitatory connection from the right S2 to the right V1 in the somatosensory-visual condition. Finally, somatosensory deviants were modulated by an excitatory connection from the right STG to the right S2 following auditory standards, and from the right V5 to the right S2 following visual standards. These results demonstrate that while the core processing of a deviant stimulus relies on its own sensory hierarchy, the network is flexibly reconfigured to incorporate information from the modality in which predictions were formed.

**Figure 5:**
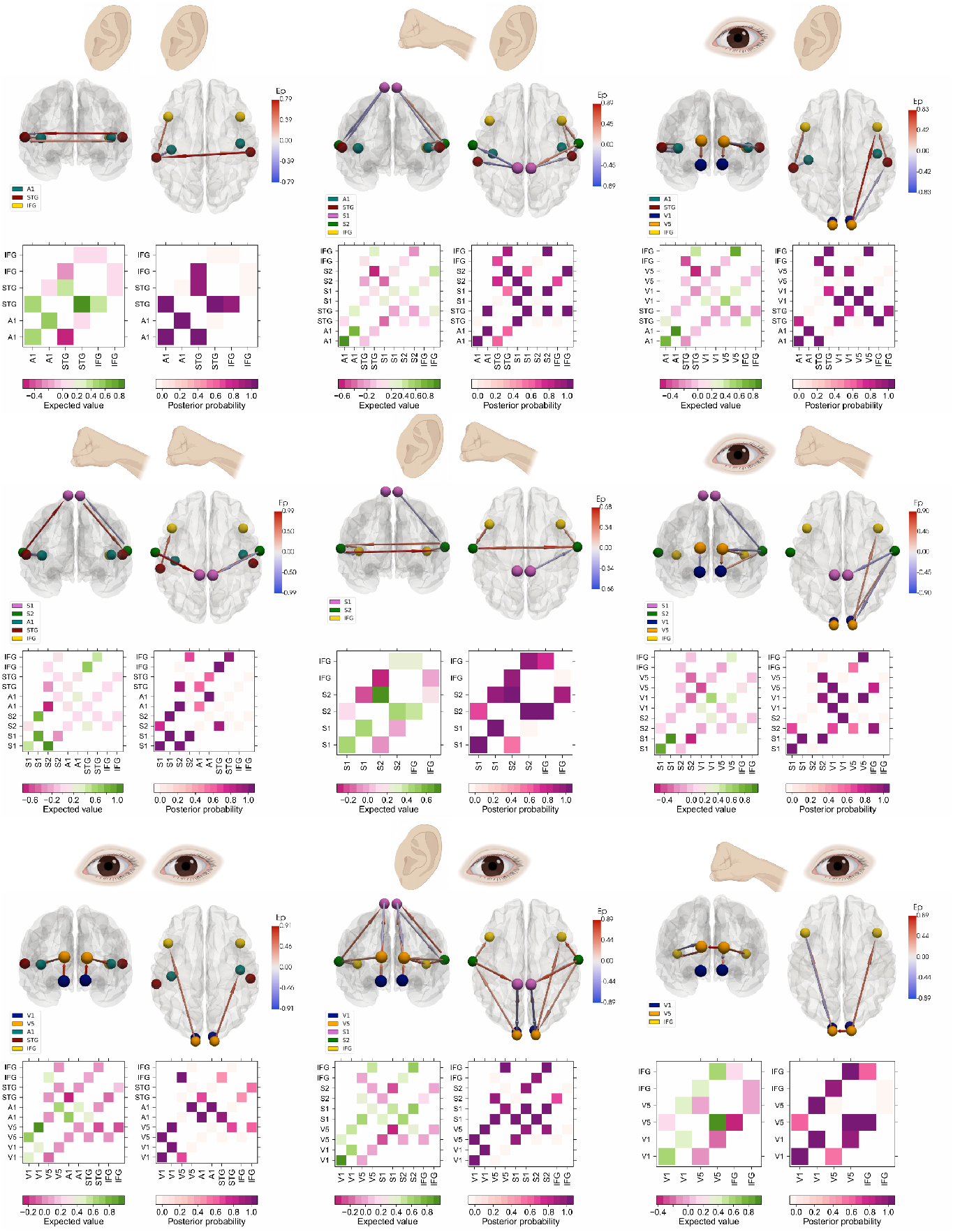
Effective connectivity networks supporting crossmodal and unimodal contrasts. This figure shows effective connectivity networks supporting crossmodal and unimodal contrasts modelled using PEB. Each glass brain panel represents connectivity differences between these conditions shown from superior and occipital view. Extrinsic connections between cortical regions are shown as arrows, with their color gradient (red-to-blue) indicating the posterior expectation of connectivity parameters, calculated from a multivariate normal probability density over PEB parameters. For clarity, only connections with a posterior probability greater than 95% (representing very strong evidence based on free energy) are included. The lower section of the figure includes two heatmaps: the left displays the posterior expected connectivity parameters, and the right shows their corresponding posterior probabilities. Sensory stimulation modalities are represented by symbols (ear = auditory, eye = visual, hand = somatosensory).

To identify how prediction error responses modulate effective connectivity differently depending on the sensory context, we performed DCM and PEB modelling for six unimodal–crossmodal contrasts shown in Figure 6. Crucially, all contrasts isolated the deviant modality while manipulating whether standards were from the same or a different sensory modality. This served to isolate crossmodal context effects on effective connectivity within dedicated sensory hierarchies. Across contrasts, we discovered that unimodal and crossmodal prediction errors consistently modulated connectivity within the same cortical hierarchy but modulations followed distinct patterns. We found that differences between uni-and crossmodal effects varied across levels and hemispheres, revealing diverse modes of crossmodal context-sensitive routing for prediction errors.

**Figure 6:**
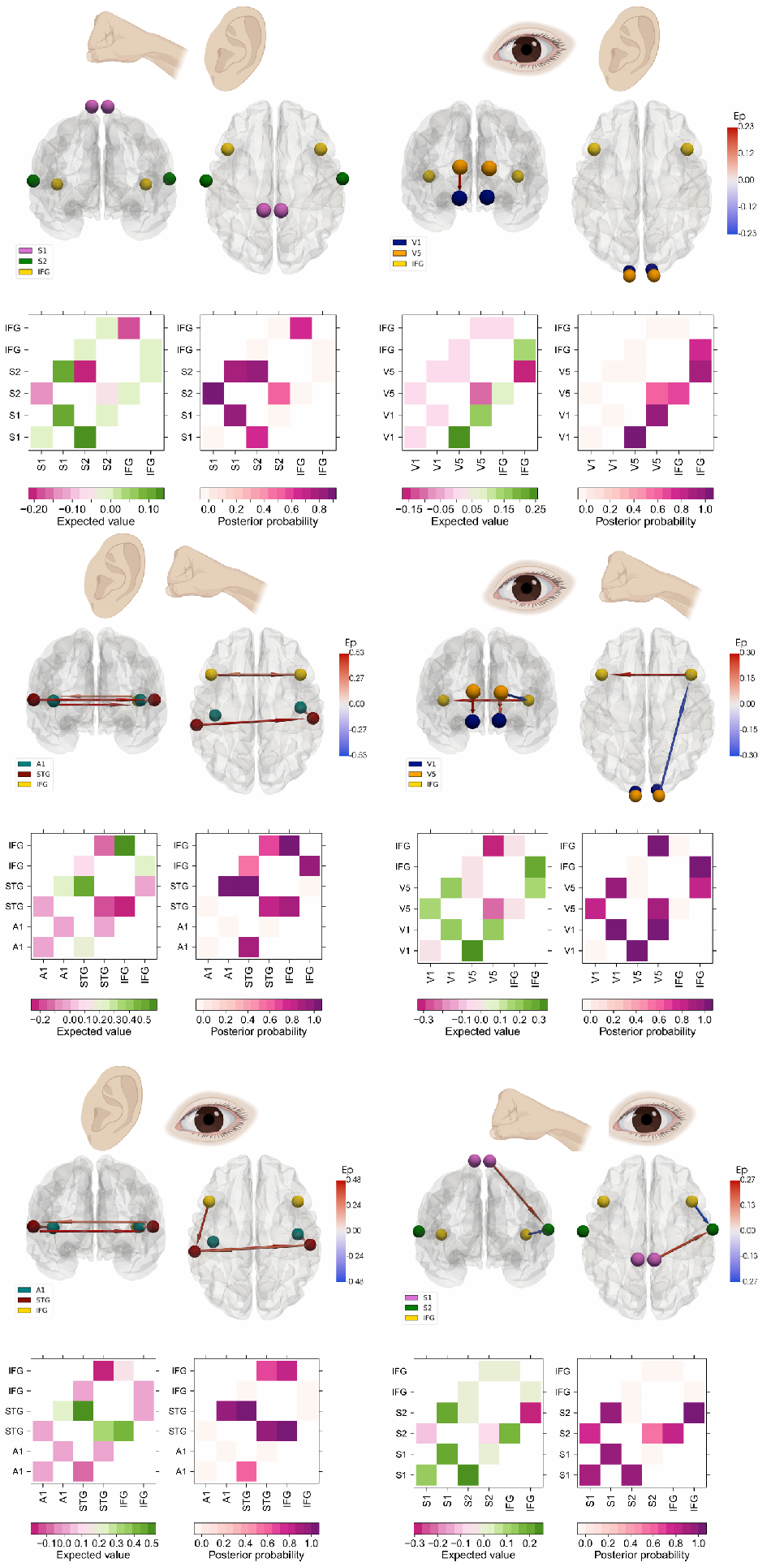
Connectivity differences between unimodal and crossmodal contrasts. Each panel illustrates connectivity differences between deviant and standard trials for each contrast modelled using PEB. Arrows within the glass brains shown in occipital and superior view depict forward and backward extrinsic connections between cortical regions. Their color (red-to-blue gradient) reflects the posterior expectation of connectivity parameters derived from a multivariate normal probability density over PEB parameters. Only connections with a posterior probability exceeding 95% (indicating very strong evidence based on free energy) are displayed. Below the glass brains, a heatmap on the left shows the posterior expected connectivity parameters, while the corresponding posterior probabilities are presented on the right heatmap. Sensory stimulation modalities are indicated by symbols (ear = auditory, hand = somatosensory, eye = visual).

#### Visual-somatosensory vs. somatosensory

When comparing prediction error responses to somatosensory deviants embedded in visual as compared to somatosensory standard streams, we identified changes in connectivity only in a right-lateralised somatosensory network. We observed increased excitatory connectivity from the right S1 to S2, and a reduction in top-down inhibition from the right IFG to S2 in the visual-somatosensory relative to the somatosensory condition. This suggests enhanced right-lateralised local integration and diminished frontal constraint in crossmodal relative to unimodal predictive coding.

#### Auditory-somatosensory vs. somatosensory

Unlike comparisons between visual-somatosensory and somatosensory prediction errors, no differences survived thresholding at a posterior probability of.95 when comparing auditory-somatosensory and somatosensory prediction errors are contrasted. However, there is strong evidence (with a posterior probability.75-.94) for an inhibition from the left S1 to the right S2, via the left S2. There was also strong evidence for excitatory connectivity between the right S1 and S2.

#### Somatosensory-visual vs. visual

The visual hierarchy exhibited stronger reciprocal interactions between right V1 and V5 in the somatosensory-visual relative to the visual condition. Moreover, frontal interhemispheric connectivity from left to right IFG was increased. Crossmodal prediction errors were supported by inhibitory connectivity from right V5 to the right IFG. We found reciprocal increased connectivity between the right S1 and S2, and from left S1 to S2 for somatosensory-visual relative to visual contrasts.

#### Auditory-visual vs. visual

In contrast to the distributed effects above, auditory-visual prediction errors affected only early visual stages. We observed increased excitation from left V5 to V1, suggesting early-stage reconfiguration without broader frontal or bilateral effects.

#### Visual-auditory vs. auditory

Crossmodal prediction errors enhanced connectivity from right A1 to ipsilateral STG, as well as bidirectional coupling between STGs. We also found excitatory connectivity from the left IFG to ipsilateral STG. This suggests that crossmodal inputs in this contrast recruited both intra- and interhemispheric temporal regions and engaged frontal nodes more strongly than unimodal auditory deviants.

#### Somatosensory-auditory vs. auditory

Unlike the visual-auditory to auditory contrast, comparisons of somatosensory-auditory and auditory prediction errors increased bidirectional IFG–IFG interactions and enhanced connectivity from right A1 to ipsilateral STG. A similar increase in interhemispheric coupling from the left to the right STG was also observed. This suggests that somatosensory-auditory prediction errors recruited both intra- and interhemispheric temporal regions and engaged frontal nodes more strongly than purely auditory prediction errors.

Taken together, information flow in modality-specific sensory hierarchies distinguishes prediction error signals when expectations are formed and violated in different sensory modalities as compared to the same. We show that crossmodal prediction errors reconfigure connectivity across multiple stages of processing relative to unimodal prediction errors, with specific differences depending on the involved modalities. While some connections such as feedforward links from early to mid-level sensory areas were commonly modulated across comparisons, others, especially those involving frontal and interhemispheric interactions, were specific to particular crossmodal contrasts. These results highlight that crossmodal expectations exert modality-specific influences on dedicated sensory hierarchies. This challenges models positing uniform top-down or feedback modulation and instead supporting a flexible, distributed model of predictive coding across the senses.

## 4 Discussion

Our study shows that the neural dynamics and mechanisms supporting crossmodal prediction errors differ fundamentally from those elicited within a single sensory modality. By combining temporal decoding and effective connectivity modelling, we determined that predictions formed and violated in different sensory modalities shape prediction error signals across early and late levels of the cortical hierarchy. We uncovered supramodal activation patterns for crossmodal prediction errors likely reflecting the convergence of sensory information from multiple channels for predictive coding across early and late levels of cortical processing. This way, we provide evidence that the temporal stability of prediction error representations in the cortex is shaped by whether sensory expectations and violations occur within or across modalities. However, distinct connectivity architectures supported predictions made across crossmodal rather than unimodal transitions. Where network architectures supporting crossmodal prediction errors diverged from their unimodal counterparts, they engaged the cortical hierarchy in a distinct manner for each combination of sensory modalities.

Previous work has established that crossmodal violations can generate mismatch responses in the cortex, extending predictive coding frameworks to multisensory contexts (Matusz, Retsa, and Murray 2016; Huang et al. 2024; Ghosh, Talwar, and Banerjee 2024). Our findings advance this understanding by showing that crossmodal prediction errors are characterised by more prolonged and temporally generalised cortical representations compared to unimodal prediction errors. Whereas unimodal prediction errors yielded transient, largely time-localised neural patterns in a mid-latency time window (Niedernhuber et al. 2022), crossmodal prediction errors exhibited neural activity which was maintained extensively at each stage of cortical processing. In line with the notion that the entire cortex processes multisensory information (Ghazanfar and Schroeder 2006), our findings suggest that cortical computations reconciling discrepancies between predictions and inputs spanning different sensory modalities are ubiquitous and distributed in the cortical hierarchy. Advancing beyond earlier results which point to a centro-parietal hub for audiovisual prediction errors (Huang et al. 2024), we uncovered unique combinations of information flow between primary sensory, temporal, parietal and frontal regions for different combinations of somatosensory, visual and auditory crossmodal prediction errors.

Earlier debates were concerned with whether unimodal and crossmodal processes could be differentiated by localising them at early vs late stages in the cortical hierarchy (Driver and Noesselt 2008; Faivre et al. 2018). Rather than focusing on the location of neural prediction error responses alone, we demonstrate that supramodal neural dynamics support crossmodal prediction errors across all levels of the cortical hierarchy. In alignment with the notion that local and global effects are supported by a supramodal activation pattern (Niedernhuber et al. 2022; Sanchez et al. 2020), we found evidence for common supramodal networks encoding both unimodal and crossmodal local and global effects. Although crossmodal local prediction errors are supported by supramodal activation patterns, crossmodal prediction errors reconfigure intra- and interregional interactions within sensory-specific hierarchies in a modality-dependent manner. While classical hierarchical predictive coding models often posit uniform top-down updating terminating in common hubs (Friston 2008; Rao and Ballard 1999), we observed that distinct crossmodal combinations led to selective changes at different levels in the cortical hierarchy. For example, somatosensory-visual prediction errors increased reciprocal coupling within the visual hierarchy and enhanced interhemispheric frontal interactions, whereas auditory-visual violations primarily altered early feedforward processing in visual regions. This pattern contradicts models assuming that crossmodal prediction errors are uniformly resolved by canonical multisensory integration centres (Calvert 2001; Talsma 2015), instead supporting more flexible and context-sensitive network adaptations across the cortical hierarchy. In sum, our results refine theoretical accounts of predictive processing by directly comparing uni- and crossmodal prediction errors. They support models proposing that prediction errors in multisensory contexts are resolved through distributed, modality-contingent adjustments (Talsma 2015; Cao et al. 2019). Overall, our findings indicate that the brain does not simply generalise unimodal predictive strategies to multisensory environments but implements tailored network adaptations depending on the specific combination of senses engaged.

We note several limitations for this study. To avoid tiredness, our design did not explore the influence of stimulus onset asynchronies, simultaneous multisensory stimulation, or probabilistic variations in crossmodal contingency which might further influence crossmodal prediction error architectures. Since subject-wise MRI was unavailable, the spatial accuracy of our connectivity results is limited. Our effective connectivity models focused on predefined cortical areas located within modality-specific cortical hierarchies. However, incorporating broader whole-brain network approaches could uncover additional routes of crossmodal integration. Although a strength of our paradigm is that behavioural responses are not necessary to discriminate prediction errors at successive hierarchical levels, linking these network adaptations directly to behavioural markers of prediction error resolution remains an important direction for future studies.

This study extends a line of work that began with the introduction of the local–global hierarchical oddball paradigm (Bekinschtein et al. 2009), which established that nested sensory regularities can be used to probe hierarchical predictive coding. Building on this framework, our 2022 study (Niedernhuber et al. 2022) demonstrated that unimodal prediction errors propagate along modality-specific hierarchies while converging on late supramodal responses. More recently (Niedernhuber et al. 2025), we showed that when two features within a modality are simultaneously violated, double-deviants evoke earlier and stronger supramodal responses than single-deviants, underpinned by inferior frontal connectivity together with modality-specific sensory pathways. The present work moves beyond both unimodal hierarchies and multi-feature violations to investigate prediction errors occurring across modalities when expectations generated in one sense are violated in another. We find that crossmodal and unimodal prediction errors differ: Crossmodal prediction errors are temporally sustained, supported by prolonged supramodal representations, and accompanied by flexible, context-sensitive reconfigurations of effective connectivity. Taken together, these studies trace a trajectory from the original description of hierarchical prediction error signatures, through their characterization in unimodal and multi-feature contexts, to the current demonstration of how predictive coding flexibly adapts to transitions between sensory modalities.

In summary, our results demonstrate that crossmodal prediction errors elicit temporally sustained cortical representations and induce flexible, modality-specific reconfigurations of effective connectivity across sensory hierarchies. These findings challenge simplified models of uniform predictive updating and advance hierarchical predictive coding frameworks by highlighting the distributed, context-sensitive nature of multisensory prediction error processing in the human brain.

